# Aberrant Neuronal Synchronization Associated with Cognitive Deficits in a Rodent Model of Childhood Cranial Irradiation

**DOI:** 10.1101/2024.12.19.629321

**Authors:** Sebastian A. Barrientos, Elise Konradsson, Crister Ceberg, Per Petersson

## Abstract

Cranial radiation can be a life-saving intervention in pediatric brain cancer therapy but often results in debilitating cognitive decline. To clarify the underlying mechanisms of these side-effects we have here recorded neurophysiological activity in distributed brain networks involved in decision-making and memory functions in adult rats exposed to cranial irradiation on postnatal day 21. Multi-structure local field potential (LFP) recordings revealed decreased power in irradiated animals in the 4-9 Hz frequency band. Additionally, a distinct slowing of the oscillatory activity was observed preceding erroneous choice in a decision-making task. Moreover, irradiated rats showed reduced dynamics and a fragmented pattern of inter-structural coherence across different phases of the task. Our results suggest that the cognitive deficits and reduced processing speed following irradiation of the juvenile brain arise as a consequence of changes in long-range functional connectivity, including thalamocortical circuits, causing abnormally slow and spatially fractionated patterns of coordinating LFP activity.

## Introduction

Cognitive dysfunction is a common sequela of brain tumor therapy in early life. In particular, cranial irradiation often causes debilitating cognitive decline ^1,2^, which severely limits the use of this very important and sometimes life-saving therapy in young children ^3^. The underlying mechanisms giving rise to these troublesome side effects are, however, largely unknown. In fact, cognitive symptoms often appear long after the procedure, even in patients with no obvious deviations in radiological assessments ^4^. Thus, while the structural injuries inflicted may be relatively subtle, they nevertheless may have severe consequences at the level of physiological processing of information in cognitive brain circuits ^5,6^. Brain cancer survivors have reported more severe cognitive side effects when cranial irradiation was applied at a young age than when irradiated later in life, likely due to interference with the normal brain development process ^7,8^.

However, to elucidate how cognitive brain circuits are affected by irradiation at an early age in essence requires direct measurements of the neurophysiological activity of the involved brain circuits with high spatial and temporal resolution during various types of mental processing. This has to date been technically difficult to accomplish using *in vivo* models. We here aimed to bridge this knowledge gap by exploring patterns of shared dynamics of neuronal populations in different brain areas using specially designed microelectrode arrays targeting distributed interconnected neuronal circuits ^9^.

Because oligodendrocytes constitute a major proliferating cell group in the brain during early postnatal years, it is plausible that an insult caused by ionizing radiation in this critical phase of brain development could lead to lifelong abnormal myelination of axonal nerve fibers ^10^. Even if such abnormalities are present on a minor scale, this could nevertheless disturb long-range neuronal communication between spatially separated brain areas, interfering with cooperative processing of information in larger neuronal networks, which is thought to be of particular importance for cognitive functions ^11^. Although previous neurophysiological data investigating this notion in the context of juvenile irradiation are scarce, evidence for connectivity-based pathophysiological network phenomena has been presented in other conditions involving dysfunctions in distributed brain circuits.

One such example is in Parkinson’s disease, where the initiation and execution of voluntary movements is believed to be hampered by pathophysiological processes involving abnormal functional connectivity in cortico-basal ganglia-thalamic circuits (see e.g., Moran et al. 2011, Santana et al. 2014 ^12,13^). If similar mechanisms are contributing to cognitive deficits in child cancer survivors exposed to cranial irradiation treatment it is likely that a combination of several factors determine the severity and characteristics of the functional deficits that are present later in life, including relative distributions of absorbed doses throughout different tissue volumes, relative sensitivity of the exposed cell groups as well as their function role in larger brain networks ^14^.

Thus, to clarify the neurophysiological mechanism whereby cranial irradiation early in life leads to life-long cognitive deficits, we have here studied the neurophysiological bases of working memory processes as well as context-dependent action selection in adult rats that were exposed to cranial irradiation at a developmental stage of brain maturation corresponding to three years of age in the human child (i.e. postnatal day 21 in rats; ^10^). Brain activity in several interconnected brain structures, including prefrontal cortex, basal ganglia, posterior parietal cortex and hippocampus were recorded, in parallel, in association with behavioral tests demanding attention, decision making and memory recall.

We here report that, during cognitively demanding tasks, irradiated rats differed in at least three ways to control rats. In specific, LFP power and peak frequency are decreased in irradiated animals in the 4-9 Hz frequency band, and this coordinating LFP population activity is particularly slowed down preceding erroneous choice in a decision-making task. Irradiated rats also showed reduced dynamics and a fragmented pattern of inter-structural coherence across different phases of the task. These findings suggest that the cognitive deficits and reduced processing speed that is a common sequela of juvenile brain irradiation arise as a consequence of changes in long-range functional connectivity, including thalamocortical circuits, causing abnormally slow and spatially fractionated patterns of coordinating LFP activity.

## Results

### Behavioral tests of cognitive function in rats exposed to juvenile irradiation

We initially performed a set of experiments aimed at screening for memory deficits induced by juvenile cranial irradiation. For this purpose, 10 rat pups were subjected to whole brain (WB) cranial irradiation at postnatal day 21 (+/-1 day) receiving a single dose of 10 Gy (200 kV x-rays), and 23 age-matched rats subjected to a sham procedure (involving all methodological steps except exposure to radiation) served as the control group (CTRL). After reaching adulthood, at eight weeks of age, animals were first evaluated in an open-field test to assess potential changes in spontaneous behavior. No significant differences were observed between WB and CTRL with respect to the total distance travelled (Bootstrapping test: Mean Diff = 3.03, CI = [-7.71, 7.08], p-value = 0.41; Supplementary Figure 1), or the time spent in the center of the field (Bootstrapping test: Mean Diff = 26.43, CI = [-50.80, 41.06], p-value = 0.23; Supplementary Figure 1), indicating that the irradiation procedure did not cause deficits in broader functions such as general motor ability nor induced anxiety-like behavior, which could both complicate the interpretation of the subsequent cognitive tests.

The following day, the rats were tested in a novel object recognition (NOR) task in the same arena. In this test paradigm, object recognition is reflected in a relatively longer time spent exploring a novel object compared to a familiar one being presented 10 min apart ^15,16^. In the NOR task, no significant difference was found in the discrimination ability of WB versus CTRL rats (Bootstrapping test: Mean Diff = 0.15, CI = [-0.41, 0.41], p-value = 0.47; Figure 1A). However, it was noted that only the CTRLs showed a discrimination ability significantly above chance level (Wilcoxon Signed-Rank test: CTRL-0 p-value = 0.032; WB-0 p-value = 0.25). One week later, rats were tested in an object-location recognition (OLR) task. In this paradigm, relatively longer time spent exploring a familiar object when placed in a new location signifies a functional object-spatial association memory ^17^. In the OLR task, irradiated animals were found to show significantly poorer discrimination ability than CTRLs (Bootstrapping test: Mean Diff = 0.46, CI = [-0.33, 0.36], p-value = 8.80e-03; Figure 1B). We then expanded this analysis, by comparing the discrimination ability resulting from more restricted irradiation procedures, targeting either the hippocampal formation (HF; involved directly in memory function), or corticostriatal circuits (CS; involved in motor control and motivated behavior). However, no significant differences were found between rats subjected to HF, CS or WB irradiation designs, neither in the NOR (Kruskal-Wallis p-value= 6.17e-01, H-statistic= 1.79, df=3) nor in the OLR tasks (Kruskal-Wallis p-value= 6.90e-02, H-statistic= 7.09, df=3). These results suggest that brain systems involved in memory processes and action selection likely tightly interact in behavioral tasks requiring both components, hence making it difficult to separate their individual contributions. For these reasons we chose to focus our further analyses on the WB group, representing the most severe model (for dose absorption calculations see **Supplementary** Figure 2).

**Figure 1.**
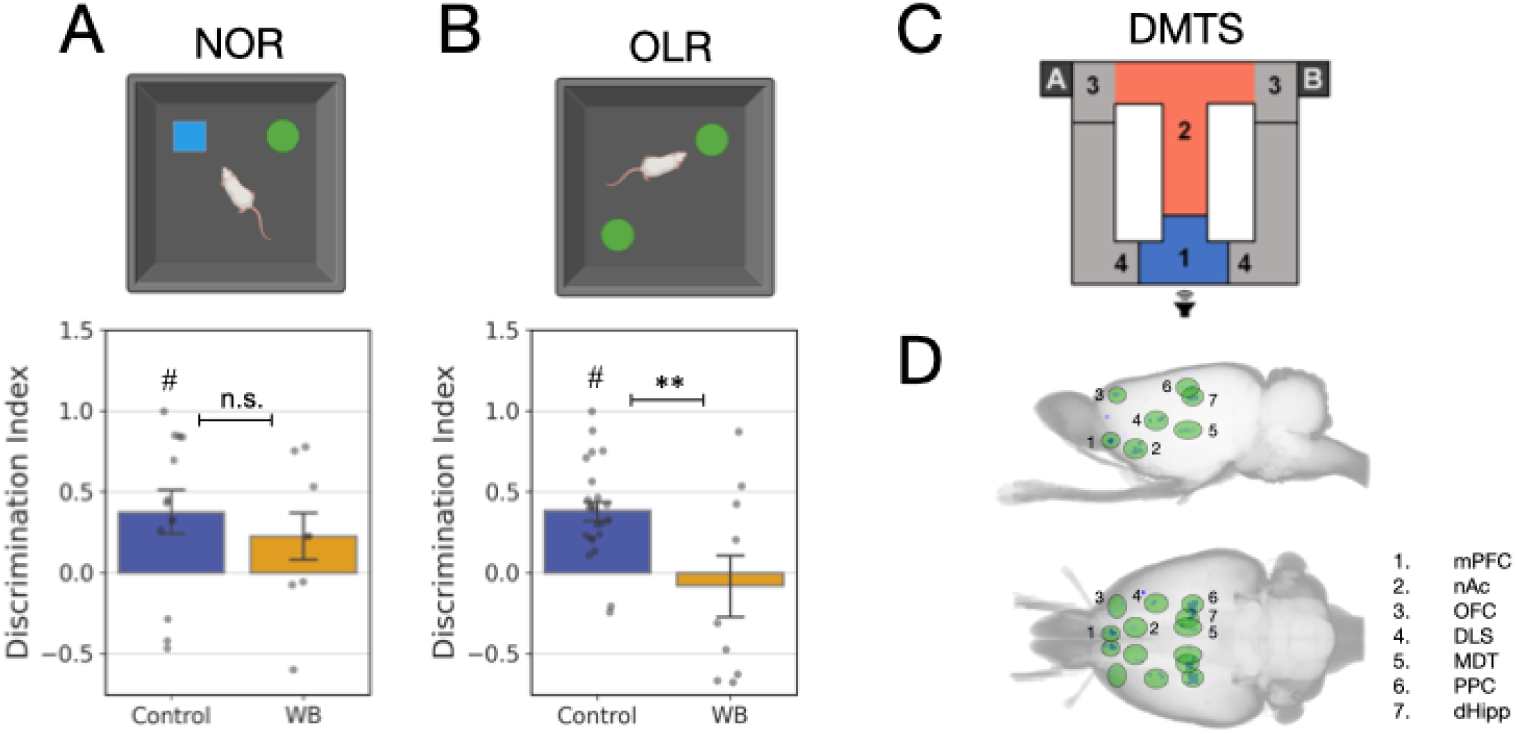
Description of behavioral tests and recording geometry. A) Top: Novel Object Recognition was performed in an open field where two different objects were presented after a habituation period. Bottom: Behavioral Performance during NOR for individual CTRL and WB rats. NCTRL-WB = [13, 8]. B) Top: Object-Location Recognition task was performed in the same setting as the NOR test but with a positioning of one of the familiar objects in a new place in the test phase. Bottom: Behavioral Performance during NOR for individual CTRL and WB rats. NCTRL-WB = [20, 11]. A-B) Bars indicate the average Discrimination Index across recorded animals and error bars indicate SEM. Filled circles indicate valid values for individual animals in each dataset. Significance was evaluated by Bootstrapping test between the samples (n.s. = non-significant, ** p-value < 0.001) and by Wilcoxon Sign-Rank test to evaluate significance against chance level (DI = 0, # p-value < 0.05). C) For electrophysiological recordings, rats were trained in a Delayed-Match-to-Sample task in an automated eight-shaped T-maze. Each trial starts at region 1 where two auditory cues are presented. When the main door opens the rat runs along the central stem (region 2) and makes the decision to turn left or right depending on if the presented cues were matching or not. Correct responses are rewarded with 0.5 mL of sucrose solution (region 3), otherwise no reward is delivered. The rat is then free to run back through region 4 to self-initiate a new trial. D) Reconstructed 3D representation based on CT scans of a representative implant showing the distribution of electrode tips spanning seven brain regions, bilaterally.

Following these initial tests, rats were next assessed in a more comprehensive behavioral test designed to probe a combination of working-memory, spatial discrimination, and action-selection in a T-maze. In this test, rats were presented with two sequential auditory cues that informed about the location of a future reward depending on if the two cues matched each other or not ( ‘delayed-match-to-sample’; **Figure 1C, top**). That is, animals had to discriminate pairs of auditory cues and retain this information in memory over a few seconds to be able to make the correct decision, turning either right (matched chirps) or left (non-matched chirps) when reaching the choice point in order to obtain a sweetened (30% sucrose) water reward. This turned out to be a very challenging task for all animals and only 60% reached the level considered to signify the learned behavior (even when using the liberal threshold of 60% correct responses as criterion). Hence, to make sure the animals being further investigated in neurophysiological recordings were indeed processing the information needed to solve the task, we elected to perform the subsequent experiments only in the sub-group of rats in each treatment group that had reached the learning criterion (i.e. at least 60% correct in 3 consecutive sessions; WB [n=5] and CTRL [n=7]). In this context it should be noted that although the selected animals in both treatment groups had learned the task and performed on a similar level, WB rats were found to be slower in their relative improvements during the learning period **(Supplementary** Figure 1). Animals were then chronically implanted with microelectrode arrays. To obtain information on differential brain activity patterns associated with different aspects of the task, we developed a specialized multi-electrode array that makes it possible to perform simultaneous recordings from up to 128 sites in widely distributed brain structures of potential importance for working memory and sensory-based action-selection. In total, electrophysiological data from 1053 recording positions, distributed bilaterally, obtained from the 12 animals, were examined (an example recording configuration reconstructed from a *post mortem* CT scan is shown in **Figure 1D**). Recording sites were then functionally grouped into seven main structures annotated according to a brain atlas for further analyses ^18^: Pre-limbic PFC (plPFC), orbitofrontal cortex (OFC), dorsolateral striatum (DLS), nucleus accumbens (nAc), posterior parietal cortex (PPC), dorsal hippocampus (dHipp) and mediodorsal thalamus (MDT).

### Neurophysiological features associated with execution of the T-maze task

Because previous studies on working memory functions in rodents have identified a crucial role of neuronal synchronization for successful task execution in similar tests ^19,20^, we focused our analyses on local field potentials (LFPs; voltage fluctuations arising from synchronized population activity of neuronal ensembles in the vicinity of each electrode ^21^). To rule out any contribution of volume conductance from sources outside the recorded structures, bipolar voltage recordings were constructed from electrode pairs located within each structure ^22^. In **Figure 2A**, an example recording of LFPs from six separate brain structures is shown during execution of a single trial in the T-maze (vertical line denotes the opening of the door). To search for candidate LFP frequency bands where population activity could serve a coordinating role between different brain structures, power density spectra were constructed for each phase of the behavioral paradigm across a frequency spectrum from 0.1 to 100 Hz. As an illustration of the dynamics in spectral content across time (with 1s-binning), an example spectrogram from nAc in a CTRL rat is shown in **Figure 2B**). Although LFP dynamics can be seen in several parts of the frequency spectrum it is evident that a band between 4-9 Hz) contains most of the dynamics during task execution. This finding was confirmed when pooling data from all animals and all trials, as indicated by the relatively higher power in this band (shaded area in **Figure 2C**). Thus, while the spectral content of the grand average of the LFPs recorded revealed no striking differences between WB and CTRL rats (**Figure 2C**; higher frequencies were also analyzed with a similar result), the power spectra clearly indicated that LFP activity in the 4-9 Hz band might be of relevance to solve the behavioral task. We therefore next specifically analyzed changes in band-power in the interval [4-9 Hz] for each of the recorded structures throughout the different phases of the task (**Figure 2D, E**). Interestingly, the band-power followed a very similar type of dynamics across the recorded structures, with power increases appearing primarily in association with the presentation of the second cue and close to the point of receiving the reward. This can be taken to suggest that working-memory processes and expectation of reward are two cognitive processes that both involve multi-structure synchronization. From this analysis it was also clear that WB and CTRL rats showed certain differences in band-power dynamics. In particular, the power increase when the animals were approaching the reward port was significantly higher in CTRLs, while the relative reduction (compared to the power outside the task) when evaluating the outcome was more pronounced in the WB group (GLME intercept: 1.87e-01, GLME interaction p-value [Treatment:RunningLate]: 0.00, df=1; Tukey HSD: Running LateCTRL-WB p-value < 1e-10; Figure 2F). Notably, the power increase during the running *per se* was not different between the groups (see early running in **Figure 2F** and **Supplementary** Figure 3), implying that the main differences in the power of LFP synchronization between the intact and irradiated brains are related to the processing of reward expectation and the subsequent trial outcome.

**Figure 2.**
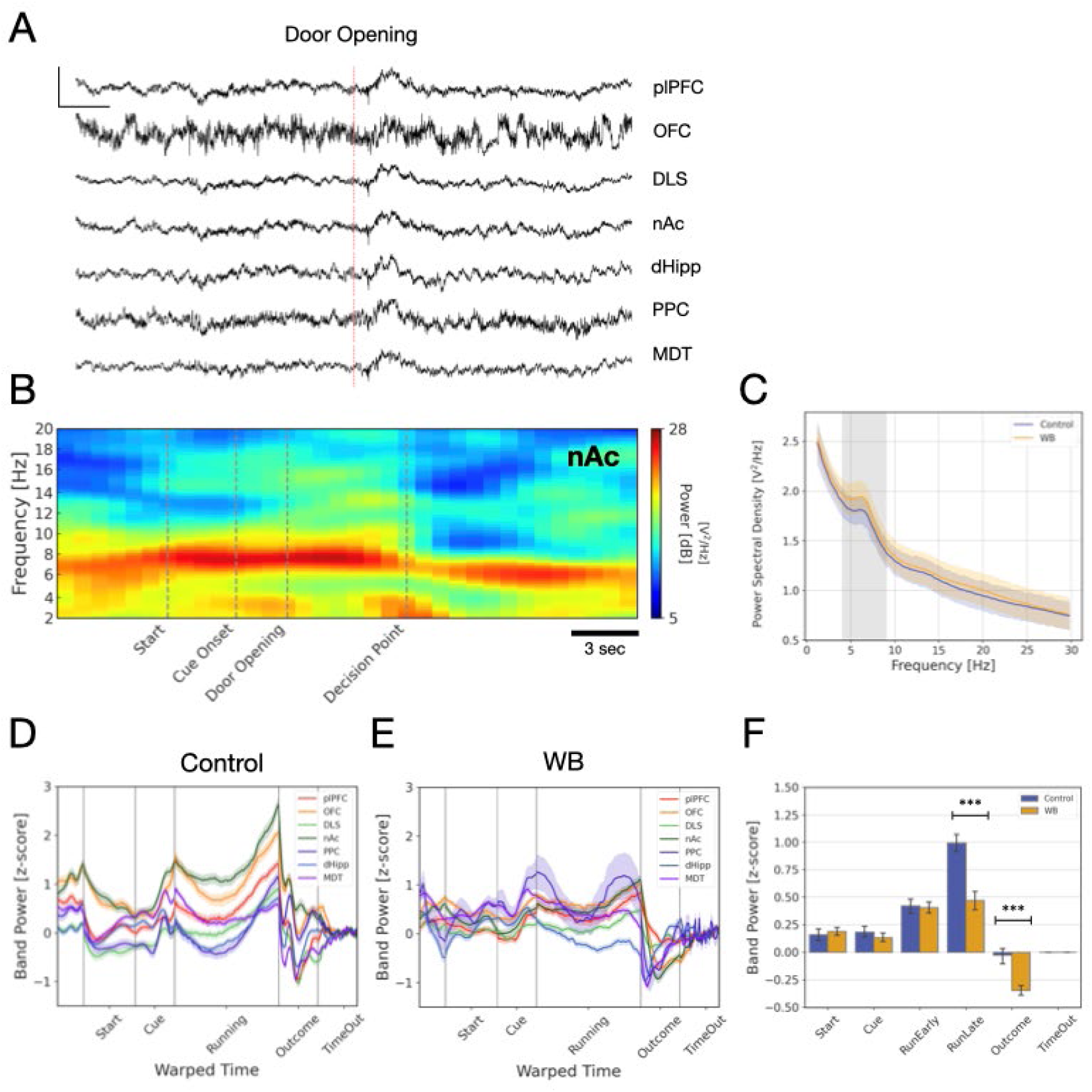
Recording of neuronal population activity in a delayed-match-to-sample test. A) Representative LFP traces centered at the opening of the main door before running through the central stem of the T-Maze. Y Scale Bar = 800 µV; X Scale Bar = 500 ms. B) Spectrogram of nAc activity during a representative trial. Vertical dashed lines indicate specific events during the trial. C) Average power spectra for all the trials recorded across all brain regions in either CTRL (blue line) or WB (yellow line) rats. The shaded area around each average line corresponds to the SEM. D-E) Normalized 4-9 Hz band power during different stages of the task for each recorded brain region in CTRL (D) or WB rats (E). F) Average 4-9 Hz band power for each phase of the task. Significant differences were found in the late running phase and during evaluation of task outcome (GLME and Tukey HSD post-hoc, *** p-value < 0.001).

### Irradiated brains show abnormal reward outcome processing

While the LFP signal consists of a great variety of different synchronized voltage fluctuations in the neuronal population close to each electrode, it has been suggested that the primary coordinating role of the LFP stems from its propensity to induce rhythmic population activity. This rhythmic activity is thought to entrain and synchronize spiking patterns across brain regions for the duration of the oscillation ^23,24^. However, LFPs recorded in freely behaving animals do not resemble pure sinusoidal waves but instead typically consist of a combination of oscillatory and aperiodic activity. Thus, to single out the dominating purely rhythmic component of the LFP from aperiodic activity we applied a normalization process that suppresses non-rhythmic components and emphasizes periodic activity ^25^. After applying this normalization procedure, it was apparent that the power distribution of the oscillatory activity (here referred to as peak fractal power) over different frequencies was also largely restricted to the 4-9 Hz band throughout the different phases of the task in both CTRL and WB rats (fractal power distributions are shown in **Figure 3A**). When comparing the relative peak power of the 4-9 Hz oscillation during different trial phases (versus Time Out), a clear difference emerged between WB and CTRL rats during the outcome evaluation phase. Specifically, while the relative fractal power increase in the irradiated brains was found to be significantly smaller than for controls (GLME Intercept= 0.83, GLME [Treatment:Outcome] Effect p-value = 0.000, nControl-WB=[200, 152]; Figure 3B), both groups showed significantly increased power when transitioning from task execution to evaluation of the outcome (Tukey HSD: CTRLrunning-outcome p-value = 0.00, WBrunning-outcome p-value = 0.00; Figure 3B). A closer inspection revealed that this difference between CTRL and WB rats in oscillatory power for task execution versus outcome evaluation occurred on a global scale and was present in all the recorded brain structures (GLME Intercept= 0.93, GLME Treatment Interaction p-value= 3.85e-08, nControl-WB=[3717, 3665]; Figure 3C). Somewhat surprisingly, however, no difference in power related to trial outcome (HIT or FAIL trials) was found for either group (GLME Intercept= 0.00, GLME Outcome Effect p-value= 0.890, nControl-WB=[178, 141]; Figure 3D). This suggests that the relative increase in oscillatory LFP power, which was more pronounced in CTRLs, likely reflects processing of the reward outcome as a generally salient event independent of its hedonic value.

**Figure 3.**
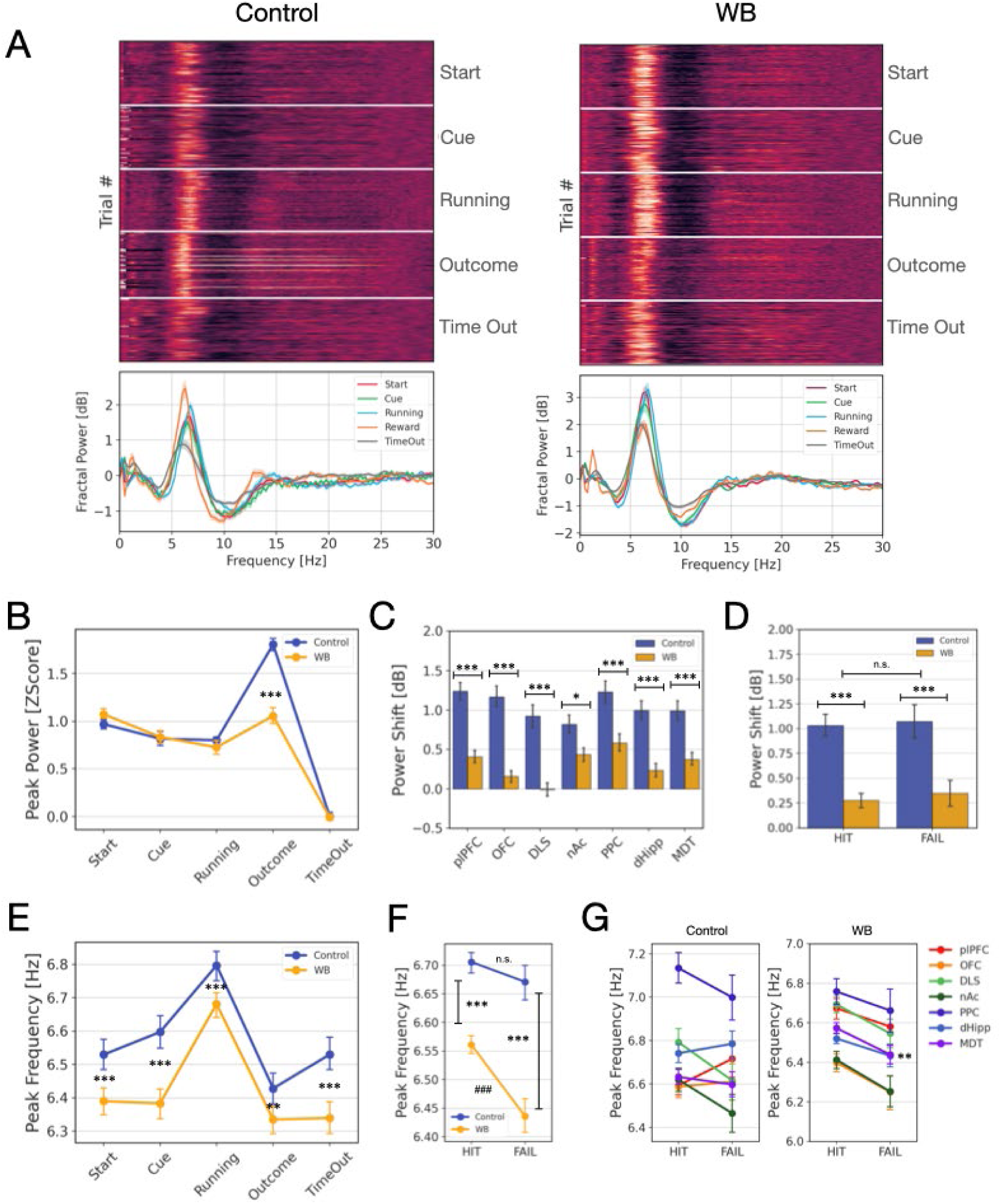
The power and frequency the power of the oscillatory LFP in the theta band differs between irradiated rats and controls. A) Power spectra normalized by 1/f displayed for each trial (top) or as the average of each stage of the task (bottom) for CTRL and WB rats. Lines indicate the mean, and the shaded area indicates SEM. B) Peak fractal power of each phase of the task normalized against the time outside the task. Replicates represent the mean of all structures per recording (NControl-WB=[388, 286]). Stars indicate a significant difference in peak power between CTRLs and WB rats. C) Comparison between CTRL and WB of the fractal power increase between the running and the outcome phase broken down into each recorded brain region. Replicates represent the mean of all trials per structure in each recording. D) Average fractal power shift for HIT and FAIL trials of each epoch compared between treatment groups (HIT: NControl-WB=[2685,2831], FAIL: NControl-WB=[1032, 834]). E) Average peak frequency of all trials of fractal spectra on each phase of the task (NControl-WB=[4081, 3726]). Asterisks mark the significant difference between treatment groups across all task phases. F) Comparison of global average peak frequency of all recorded trials of the oscillatory LFP during memory and attention demanding phases of the trial (Cue and Running phase, respectively) on HIT and FAIL trials (N HITControl-WB=[5904, 5732], N FAILControl-WB=[2188, 1718]). Note that WB rats showed a significant slowing of the peak frequency on erroneous trials (marked by ###: p-value= 0.0003). C-F) Significance was evaluated by GLME with Tukey HSD post-test (* p-value < 0.05, ** p-value < 0.001, *** p-value < 0.001). G) Average peak frequency for HIT/FAIL trials broken down into each recorded brain region for CTRL and WB rats. Error bars indicate SEM. Significance was evaluated by Mann-Whitney U –test (** p-value < 0.01, N WB-MDTHIT-FAIL= [884, 262]).

### Irradiated brains show slower population synchronization

In the analyses of peak fractal power, we noted that the peak frequency of the oscillatory component sometimes varied within the 4-9 Hz band between individual trials. We therefore next analyzed if any systematic differences in the peak frequency of the LFP oscillations, across brain structures, could be detected between treatment groups. Intriguingly, the WB rats were found to display slower oscillations than CTRLs in all phases of the task (GLME Intercept: 6.60, GLME Treatment Interaction p-value: 3.72e-04; Tukey HSD; Figure 3E). This observation led us to hypothesize that cognitive deficits after juvenile irradiation could result from a slower rhythm in neuronal population synchronization, since this oscillatory frequency would effectively limit the speed whereby coordinated computations can be performed. To directly test if the frequency of the oscillatory LFP could influence cognitive function, the fractal peak frequency during the phases of the task that required attention and involved working-memory processes (i.e. start, cue and running) was analyzed for HIT and FAIL trials separately. This comparison revealed that whereas the CTRL rats showed no differences in fractal peak frequency, the WB rats displayed a significant slowing on the trials where they made erroneous decisions (GLME Intercept: 6.71, GLME Treatment Interaction p-value: 0.00; Tukey HSD: CTRLHIT-FAIL p-value = 0.34, WBHIT-FAIL p-value = 3e-4; Figure 3F). When analyzing each of the recorded structures separately, a similar trend was observed in several structures in irradiated animals, but only MDT was found to have a significantly slower synchronization frequency on erroneous trials (p=0.0084, Bonferroni-corrected t-test; Figure 3G). Given the critical role of the thalamus in the control of global cortical dynamics, it would be a plausible scenario that this thalamic slowing could have widespread effects across various cognitive systems ^26^. Specifically, slower processing within corticothalamic and cortico-cortical circuits may contribute to impaired decision-making and delays in learning under certain conditions (cf. **Supplementary** Figure 1E-G).

Taken together, our results suggest that even though several of the irradiated animals learned the T-maze task, they may nevertheless have a double disadvantage compared to controls: First, an abnormal processing of reward outcome (as indicated by the smaller increase in oscillatory power in association with the reward phase - potentially affecting appropriate credit assignment; **Figure 3B-D**), and second, a reduced computational speed in the phases of the task demanding attention and working memory (slower neuronal population synchronization; **Figure 3E-G**).

### Irradiated brains show reduced multi-structure coordination

The discovery that irradiated brains in certain respects display deviant patterns of LFP activity made us next pose the question if these abnormalities could influence multi-structure functional connectivity. The size of the involved constellation of networks involved could be of particular importance in situations demanding higher cognitive load, since such processes are thought to rely heavily on the coordination of neuronal activity in widely distributed brain networks ^11^. Characterizing brain-wide activity patterns is obviously very experimentally challenging, but thanks to the specially designed implants used in this study, seven brain structures could be recorded, in parallel, providing an estimate of inter-structural functional connectivity based on the degree of coherent oscillatory activity in the 4-9 Hz band throughout the different phases of the task. These analyses revealed that the magnitude of the overall coherence (analyzed for each hemisphere separately) showed distinct dynamics across the task phases in CTRLs but not in the WB rats (Figure 4A). In specific, the CTRL animals had significantly increased coherence during the running phase, which was not observed in WB rats (GLME Intercept: 0.39, GLME Treatment Interaction during Running p-value: 0.00, NCTRL-WB = [534, 332]; Figure 4A-B). Interestingly, a significant difference between the treatment groups was also found in the outcome evaluation phase, specifically for successful trials (GLME Intercept: 0.39, GLME Treatment Interaction during HIT p-value: 0.00, NCTRL-WB = [534, 332]; Figure 4B). In fact, in the WB group no significant changes in coherence were observed during any of the different phases of the task for either HIT or FAIL trials (p>0.05, GLME and Tukey HSD posttest; Figure 4B).

**Figure 4.**
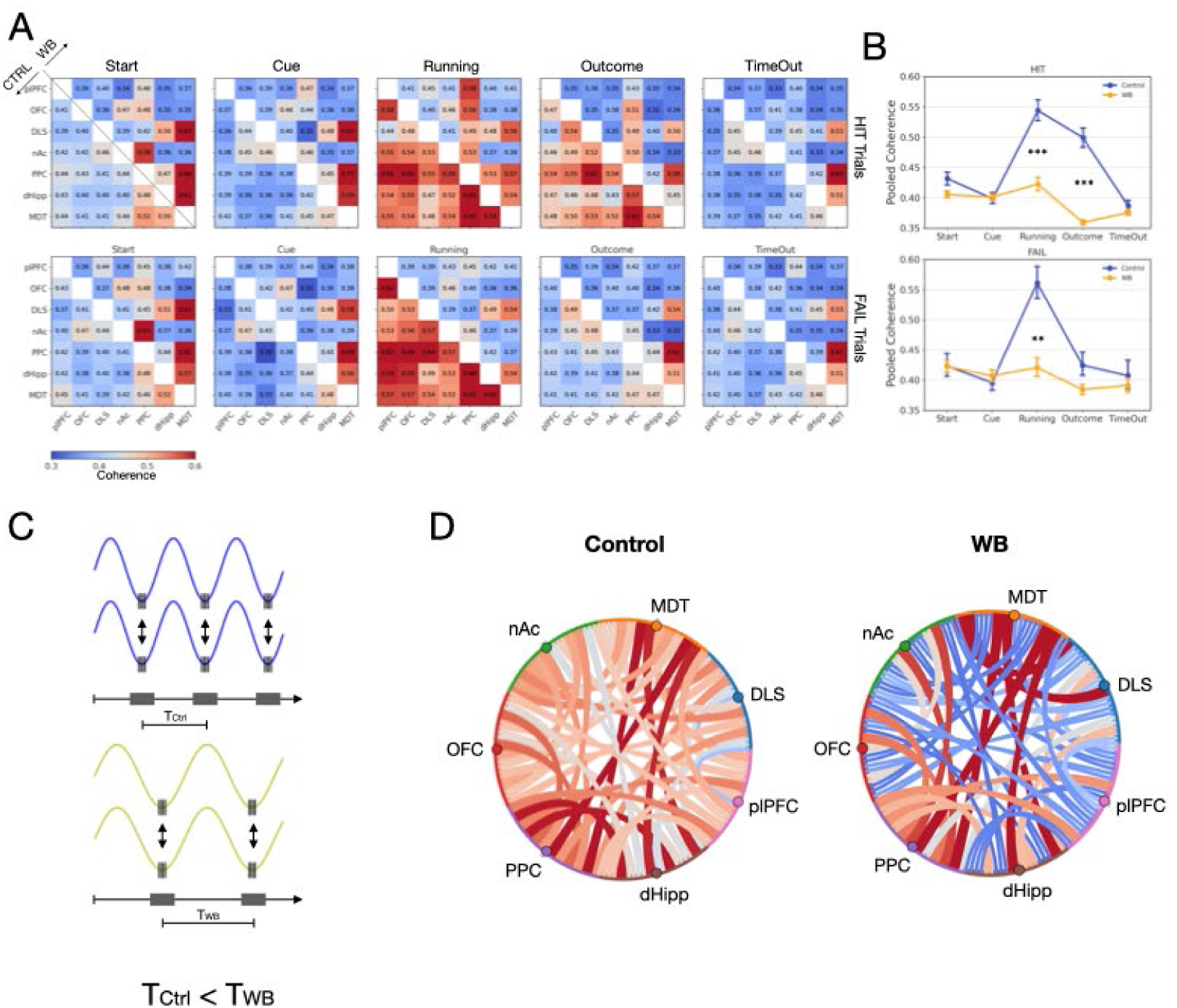
Functional connectivity in the irradiated brain shows an altered pattern. A) Pair-wise coherence in the 4-9 Hz band between each recorded brain region for HIT (top row) and FAIL (bottom row) trials. Each matrix represents a specific phase of the task. Values over and under the diagonal correspond to WB and CTRL rats, respectively. Values represent the average coherence of all trials per Outcome between the respective pair of structures for the indicated treatment and phase of the task. B) Average pooled coherence for all trials recorded for HIT (top) or FAIL (bottom) conditions. Significance was evaluated by GLME with Tukey HSD post-test (*** p-value < 0.001). C) Conceptual image illustrating principle for how synchronized population activity may control temporal windows of mutual communication. D) Chord diagram on a representative two representative recordings, CTRL and WB, respectively, illustrating the magnitude of coherence between the recorded brain regions.

Further examining the pairwise coherence observed between all pairs of structures, it became clear that WBs differed from CTRLs not only with respect to changes in the average magnitude of the coherence across phases but also in the more detailed pattern of inter-structural coordination. That is, while coherent oscillations appeared to connect all pairs of recorded structures relatively evenly in the CTRLs, this systems-level coordination tended to be much more fractioned in the irradiated brains. Instead, coherence between a few restricted structures dominated in WB rats, potentially creating islands of localized processing within the otherwise much larger global workspace being coordinated in intact brains (in quantitative terms, this was reflected as a significant difference in the coefficient of variation [CV] of coherence between pairwise structures for the different phases; average CVs were (CTRL/WB): 0.12/0.19, p<0.001; Mann-Whitney U-test). Somewhat unexpectedly, the pairs of structures with a relatively higher coherence in the WBs, did not seem to specifically include anatomically closely connected regions (Figure 4A). It can be speculated, however, that given that MDT was overrepresented among the coherent pairs it is conceivable that thalamic synchronized oscillations may be a key driver of the patchy networks of coordinated activity observed in the irradiated brains.

## Discussion

In this study we sought to clarify how exposure to cranial irradiation at a young age can lead to cognitive deficits later in life by characterizing brain activation patterns during cognitively demanding tasks in rats exposed to ionizing radiation at a juvenile age. We found that brain activity patterns differed in irradiated rats compared to controls in at least three ways. First, the band power [4-9 Hz] of the LFP ^20^ differed during the phases of the task involving processing of expected reward and task outcome (i.e. approaching and checking the reward port). Interestingly, previous studies in rats performing a similar T-maze task have suggested that synchronous LFP oscillations in the 7-14 Hz range have an important role in binding together complementary learning and memory systems – specifically, hippocampal dependent episodic memories and cortico-basal ganglia dependent systems for procedural learning and memory ^19^. Thus, it is possible that the relatively lower LFP power observed in irradiated rats during these task phases signify a reduced engagement of hippocampal memory systems (see also dHipp in **Figure 2E**). This observation would be in line with previous findings in rodent brain slice studies indicating that cranial irradiation can diminish hippocampal neuron excitability and plasticity ^27,28^. However, direct *in vivo* neurophysiological confirmation of persistent effects into adulthood has so far been lacking. The notion that the irradiated brains show a deviating processing of reward outcome was further supported by the differences observed in the power of the oscillatory component of the LFPs - which is thought to be particularly important to help coordinating neuronal activity across different brain structures ^23,24^. Indeed, WB rats displayed a relatively reduced power increase of the oscillatory LFP in the 4-9 Hz band in the reward phase on a global level.

Second, the peak frequency of the rhythmic coordinating LFP activity differed significantly between intact and irradiated animals. To our knowledge this is the first time this phenomenon has been reported and we can only theorize about its functional consequences. It seems plausible, however, that the frequency of the coordinating LFP will principally determine the processing speed of the involved neuronal networks by setting the cycle time between the shared spiking events (i.e. the time-window of the cycle when neuronal populations are in their depolarized phase facilitating mutual information transfer; **Figure 4C**), limiting the capacity of the irradiated brain ^29–31^.

In any case, the observation that a slowing of oscillatory LFP in WB rats, in the phases of the task that require attention and involves working-memory processes, was more pronounced in conjunction with erroneous trials clearly suggests that the reduced oscillatory LFP frequency has negative functional consequences. The fact that this slowing was particularly pronounced for thalamus could also explain the broad consequences for cortical networks.

Third, irradiated animals showed reduced functional connectivity between brain structures, as evident by measurements of the inter-structural coherence ^12,13^. Owing to the specially designed recording arrays developed for this study, it was possible to analyze multi-structure coordination during the execution of the task on a larger scale than what has previously been achieved. This revealed several interesting differences between WB and CTRL rats. Not only did the irradiated animals show a deviant pattern in overall coherence during the different task phases compared to controls, but there was also a striking reduction in the spatial scale of the multi-structure coordination. We are not aware of previous reports describing findings on a similar temporal scale (milliseconds), but in the human fMRI literature it has been suggested that the coordination of large-scale networks, on a seconds-scale, is of particular importance to support cognitive functions ^11^. Thus, to the extent that coordinated synchronized LFP activity is supporting the cognitive processes required to make successful choices in the delayed-match-to-sample task used in the current study, it is reasonable to assume that the fragmented connectivity map observed in the WB rats (as illustrated in **Figure 4D**) entail certain disadvantages. Again, it is interesting to note that the aberrant connectivity pattern in the irradiated brain appears to be strongly connected to the thalamocortical network.

Taken together, while all rats selected for recordings were able to perform the task (which was a requirement to ensure we were comparing task-relevant brain activity), the discovered differences in power, frequency and scale of coordinating LFP activity could still have functional consequences in some situations (e.g. possibly contributing to slower improvements in WB rats during the pre-recording training sessions; **Supplementary** Figure 1E-G). However, while our findings strongly suggest that altered LFP oscillations are a key neurophysiological factor underlying radiation-induced cognitive dysfunction (possibly due to radiation damages to dividing oligodendrocytes at an early age, causing long-term abnormalities in axonal myelination) this remains a hypothesis at this point, which clearly requires further investigation. Nevertheless, if corroborated, these insights could significantly influence clinical practice. For instance, our results highlight the importance of understanding the interplay between hippocampal memory systems and action-selection networks for the development of radiation protocols that minimize cognitive impairment and point to disturbances in thalamocortical coordination as a critical factor for cognitive dysfunctions. In this context, alternative radiation approaches such as ultra-high dose-rate irradiation, also known as FLASH irradiation, would clearly be of interest to evaluate, given the indications that this procedure spares cognitive functions better than conventional radiotherapy ^32^. In any case, clarifying the neurophysiological bases for the cognitive dysfunctions that arise as a consequence of this necessary and sometimes life-saving intervention is an important first step towards improving the quality of life for pediatric cancer survivors.

## Methods

### Animals

Experiments were performed in Sprague–Dawley rats (Taconic, Denmark) at postnatal day 21 (+/- 1 day), and therefore they were housed together with their mother until the experimental procedures, after which the pups were weaned. Rats were kept until adulthood for further testing housed in standard cages under controlled temperature (22 °C) and humidity (50%) laboratory conditions on a 12:12 hours light/dark cycle and had access to food and water *ad libitum*. All procedures were approved by the Malmö-Lund Ethical Committee on Animal Research.

### Irradiation Procedure

Rats at postnatal day 21 were subjected to cranial irradiation, for which the animals were anesthetized with a mixture of fentanyl/medetomidine (0.21/0.21 mg/kg, i.p.; Apoteket AB, Sweden) and ophthalmic ointment was applied to protect their eyes from drought. As a control, age-matched litter mates were subjected to the sham procedure, for which they were only anesthetized for the time the irradiation procedure lasted. Each animal was fixed in prone position by a stereotactic system and then placed individually in a custom-made ventilated plexiglass chamber equipped with an FPP-2 air filter to be transported to the irradiation facility. The treatments were carried out using 200 kV X-Rays (XStrahl, UK), and the beam was collimated to the targeted brain areas using lead apertures placed on the ventilated chamber (Supplementary Figure 2). After the procedure, animals were transported back to the animal facility and woken up with atipamezole hydrochloride (0.5 mg/kg, i.p.; Apoteket AB, Sweden). After full recovery rat pups were placed back in their home cage.

### Behavioral tests of intermediate-term memory function

#### Open Field Test

Animals between 8-12 weeks of age after irradiation or sham procedure were subjected to different behavioral paradigms. Firstly, rats were handled for 10 minutes for 5 days to reduce stress related to the manipulation by the experimenter (Bevins & Besheer, 2006). After each handling, rats were allowed to explore an Open Field (OF) arena of 50x50 cm for 5 minutes. After this period of habituation, rats were video recorded exploring the arena for 20 minutes from the ceiling of the setup. Metrics of locomotion activity were extracted offline through automatic animal tracking from the videos. The distance traveled was calculated as the difference of the position of the animal centroid from t_n_ to t_n+1_. To evaluate anxiety-like behavior, center occupancy was calculated as the number of video frames, within the inner 50 percent of the OF.

#### Novel Object Recognition

Twenty-four hours after the OF-test, the rats were subjected to a Novel Object Recognition (NOR) test ^33,34^. In the same arena rats were habituated, animals were allowed to explore for 15 minutes two identical objects placed in two different corners. Rats were then returned to their home cage for 10 minutes ^15,16^, after which the rats were brought back to the arena for 5 minutes for the test phase, in which one of the two objects was exchanged for a new one in its corresponding place. Object visits were detected offline as all the times the nose of the animal was within a 10 cm radius from the center of the object. Memory performance was assessed using the difference of the time spent exploring the novel object versus the familiar object relative to the total time (first 2 minutes of the test phase). This metric is referred herein as the discrimination index ^17^

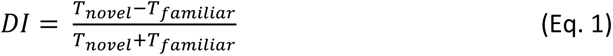

#### Object Location Recognition

Similar to NOR, the animals were allowed to explore two identical objects placed in the arena for 15 minutes. After 10 minutes pause in their home cage, the animals were reintroduced on the arena for 5 minutes for the test phase, in which one of the familiar objects had been relocated to a new position ^17,34^. Memory performance was evaluated using the DI within the first 2 minutes of the test phase (Equation 1).

### Video Tracking Analysis

Behavioral tests were video recorded, and the position of animals was tracked offline with the open-source tool DeepLabCut ^35^. A total of 600 frames picked randomly from 50 videos with different brightness conditions were used to label different body parts of the rat in order to train a deep learning network over 1030000 iterations. The network was then fed with all top camera videos to track the chosen body parts. Next, x and y coordinates of different body parts, i.e. nose tip, left and right ear, body center (point on the midline running on the back between the thorax and abdomen), base of tail, and tip of tail, were extracted using MATLAB (MathWorks), considering only a detection likelihood of > 0.9. The centroid of the animal was calculated by averaging the x and y coordinates of all 7 body parts.

### Cued Delayed Match-to-sample Task

21 rats were trained to self-initiate trials and to recognize whether pairs of auditory cues matched or not to identify in which arm of an 8-shaped T-maze (Maze Engineers) a liquid reward was available. For this, animals were restricted of water for 24 hours and habituated to explore the maze and it could identify the location of the reward (0.2 mL of 30% sucrose solution) in both arms of the maze. Next, the rats were subjected to 14 training sessions and every 5 days rats were allowed to rest from the water restriction for 2 days. In each training session a rat was placed at the start area with the doors closed, where a 5-second-long white noise was presented, indicating the start of the trial followed by a two 1-second-long auditory cues separated by a 0.5 second. Matching chirps (ascending or descending between 0.5 kHz to 5 kHz) indicated that the right arm was baited while mismatching chirps indicated the opposite arm. After a variable delay between 0.3 and 2 seconds, a 0.1 second click was played indicating the opening of the door to the central stem of the maze 0.5 seconds after. After the rat made its decision and entered to one of the side arms an infrared sensor triggered the close of the opposite arm and the release of the reward only in case of a correct response. The animal was then free to go back to the start position and initiate a new trial. This cued ‘delayed-match-to-sample’ test was designed to probe working memory capacity and executive functions (note, however, that the spatial component of the task being performed in a familiar context can be taken to suggest the additional involvement of e.g. hippocampal circuits and related structures; **Figure 1E**; ^36^). Each event in the task was controlled by a custom-made script on C# to coordinate the readings of the IR sensors of the T-Maze with the timing and delivery of specific actions (i.e., opening and closing doors and delivering reward). Task events were synchronized with an Open Ephys recording system and analyzed offline.

### Criteria for distinguishing learned behavior in T-maze

Only a certain fraction of animals in all experimental groups showed gradual improvement until reaching a steady learning criterion of 60% of correct responses for at least 3 sessions, at least 10 completed trials, and a significant, positive linear regression between the performance of the first and the second week. Rats reaching this criterion are referred herein as ‘learners’, as opposed to the remaining group of non-learners, which were excluded from further neurophysiological studies.

### Electrode implants

Recording implants of 128 insulated tungsten wires (33 µm diameter; California Fine Wire Company, CA, USA) were built as previously described ^9^. The microwires were arranged into 8 electrode bundles per hemisphere with 250 µm spacing between wires in each horizontal dimension and a wire length corresponding to the depth of each recording target. A 125-µm silver wire (Advent Research Materials Ltd, UK) was used for ground connection. All wires from each hemisphere were connected to a custom-designed printed circuit board with silver conductive paint (Electrolube, UK) and secured with UV-curable adhesive (Dymax, CT, USA).

Electrode implantation surgeries were performed after completion of the training period and reaching the learning criterion. For this, animals were anaesthetized with fentanyl/medetomidine (0.21/0.21 mg/kg, i.p.; Apoteket AB, Sweden) and fixed to a stereotaxic frame (David Kopf Instruments, CA, USA) in a flat skull position. Craniotomies were drilled over the target recording sites followed and dura was cut and removed to access the cortex. Silver wire was attached to three screws in the occipital skull bone for ground connection. The implant was anchored to the skull screws with dental acrylic cement (Kerr, CA, USA) also covering the ground wires for electrical insulation. After surgery, the anaesthesia was reversed by atipamezole hydrochloride (0.5 mg/kg, i.p.; Apoteket AB, Sweden), and buprenorphine (0.05 mg/kg, s.c.; Apoteket AB, Sweden) was administered as postoperative analgesic. The animals were allowed to recover for a minimum of 1 week before recording sessions commenced. During this period, they received extra postoperative care until fully recovered.

### Electrophysiological Recordings

Neural activity was acquired using an Open Ephys system with four Intan RHD2132 amplifier chips (Intan Technologies). Wideband signals (1 Hz - 10 kHz) were digitized at 30 kHz. Local field potentials (LFPs) were extracted offline, low-pass filtered at 500 Hz (8th order Butterworth), and downsampled to 2 kHz.

Bipolar LFPs were derived from electrode pairs within each bundle to minimize contributions from non-local sources. Time-frequency power spectral densities (PSD) were calculated over the 0–100 Hz frequency range for the whole duration of each phase of the task on each trial. The time average PSDs were then warped by averaging the power within a moving window of 0.5 Hz with no overlap.

To accentuate oscillatory components, a 1/f normalization procedure was applied ^25^. This involved fitting a power-law function (1/f^α) to the PSD and then dividing the original PSD by this fitted function. This normalization effectively removes the aperiodic, or “background,” component of the spectrum, which typically exhibits a 1/f-like distribution. The resulting normalized power spectrum emphasizes the periodic, or oscillatory, components, making them more prominent for subsequent analysis. Fractal-normalized power was calculated across the entire period between selected events, mirroring the PSD approach.

### Motor Behavior

To evaluate the magnitude of motor behavior during the electrophysiological recordings, signals from head-mounted accelerometers were recorded at a sampling rate of 30 kHz using analogue 3-axis accelerometers (ADXL335), in-built in the recording headstages (Intan Technologies). Accelerometer data was extracted offline, downsampled to 2000 Hz and followed the same treatment as LFPs. The signal was rectified, and after calculating its PSD, the signal from all accelerometers was averaged and the band within 1-45 Hz frequency range was used for further analyses.

### Functional connectivity

To evaluate functional connectivity between different brain regions, two random electrode pairs were chosen from each region, and their coherence was calculated using the *coherencyc* function from the Chronux toolbox^37^. This analysis was performed for each task phase on every trial, as described above. The average coherence within the 4-9 Hz frequency band was then used to construct a connectivity matrix, representing the strength of functional interactions between the different brain regions.

### Statistics and reproducibility

The normality assumption was assessed using the Kolmogorov–Smirnov test, and since normality was rejected, non-parametric statistics were used. For single-factor analyses, the Kruskal-Wallis test with Dunn’s post-test was applied for comparisons involving three or more groups, while the Mann-Whitney U test was used for two groups. In some cases, a bootstrapping test with 10,000 resampling iterations and 95% of confidence interval was performed, deriving the observed statistic and p-value from the resulting bootstrap distribution. For paired tests, a Wilcoxon Signed-rank test was employed. To account for the interaction of irradiation treatments and repeated measures, i.e. same animals compared in several phases of the task, analyses were conducted using a Generalized Linear Mixed-Effects model (GLME) and a Tukey-HSD post-test when interaction was significant. Comparisons with p-values ≥ 0.05 were considered not significant. Replicates in analyses involving behavioral performance were considered as the average per animal. Unless stated otherwise, brain activity measures were evaluated for each trial during electrophysiological analyses. Samples sizes are described in the main text and in the respective figure legends. Significant p-values are represented with asterisks in the order *p* < 0.05 *, *p* < 0.01 **, *p* < 0.001 ***. Error bars represent mean ± SEM.

### Characterization of dose distribution for different radiation geometries

To account for potential differences between brain structures in the characterization of the neurophysiological effects of childhood cranial radiation, we here expanded on a previously established rodent model that captures several important aspects at a molecular, morphological, physiological and behavioral level ^38–41^. In specific, we included three different radiation procedures affecting different volumes of the brain parenchyma. Each procedure, using different radiation aperture, was designed to expose primarily either the hippocampal formation and parietal association areas (PPC-Hipp), known to be directly involved in memory acquisition processes ^42^, or corticostriatal (CS) circuits, which are regarded as crucial for appropriate action selection ^43^, or alternatively a wider region spanning both sub-volumes (here referred to as whole-brain; WB).

To obtain reliable estimates of the absorbed doses in the different brain volumes, a detailed dosimetric model was constructed for each of the apertures, using the μRayStation software (RaySearch Laboratories, Sweden), which was subsequently verified by quantitative measurement using radiochromatic film in a sold water phantom (**Supplementary** Figure 2).

Based on the simulation, it was clear that sub-cortical structures such as the basal ganglia received relatively similar absorbed dose compared to different parts of the neocortex as well as the hippocampal formation. Thus, the different cell-types in the brain tissue should be affected to a relatively equal extent in both in the overlying cortex and in the deeper structures, including the basal ganglia and the hippocampal formation. Moreover, the different apertures proved to generate distinct dose distributions where the whole-brain procedure resulted in very similar doses in PPC-Hipp and CS volumes, as well as for the rest of the cerebrum, whereas the localized PPC-Hipp and CS procedures were very selective (**see Supplementary** Figure 2**, right column).**

### Computed tomography for electrode positions verification

To determine the final position of the recording electrodes in the brain, we used a semi-automated method for segmenting wires along computed tomography (CT) x-ray images of the rat heads ^44^. For this procedure, rats were anesthetized with a lethal dose of sodium pentobarbital (100 mg/kg i.p., Apoteksbolaget AB, Sweden) and subjected to transcardial perfusion of 4% paraformaldehyde. The extracted head was kept with the preserved recording implants in 4% paraformaldehyde for at least 24 hours before imaging. CT scans were conducted using a Mediso Nanoscan PET-CT scanner (Mediso, Hungary), with the head positioned such that the wires were perpendicular to the photon beam ^44^. The scanned volumes were registered to an anatomical atlas using bone landmarks and the image coordinates of the electrode tips were converted to stereotaxic atlas coordinates. Finally, the wire tips were assigned appropriate anatomical labels based on their location in the atlas ^18^.

## Code availability

All custom-written codes employed for the analyses described herein are available as Jupyter notebooks in our GitHub repository: https://github.com/NRC-Lund/radiation-WM.git

## Data availability

Research data are stored in an institutional repository and will be shared upon request to the corresponding author.

## Acknowledgements

The authors are thankful to Johan Wirén for skillful guidance in the irradiation procedures, and to Romulo Fuentes for thoughtful comments on earlier versions of the manuscript.

The study was supported by grants from Barncancerfonden (PR2018-0122), Oskarfonden, Linnea & Josef Carlsson, Sigurd och Elsa Goljes minne, Kungliga fysiografiska sällskapet, Thorsten & Elsa Segerfalks Stiftelse, Kockska Foundation, Crafoord Foundation and Kempe Foundation. Petersson also received support from the Swedish Parkinson Foundation, The Swedish Brain Foundation, Åhlén Foundation, Sven-Olof Jansons livsverk, Insamlingsstiftelserna and Vetenskapsrådet (VR) Grant 325-2011-6441, Grant 2018-02717 and Grant 2021-01769. Ceberg was supported by Mrs Berta Kamprad Foundation FBKS-2024-23 - (608) and The Swedish Cancer Society 20 1298 Pj 01 H. The computations were enabled by resources provided by the Swedish National Infrastructure for Computing (SNIC) at LUNARC partially funded by the Swedish Research Council through grant agreement no. 2016-07213.

## Authors contribution

S.A.B. designed and executed experiments, analyzed and interpreted data, developed figures, and contributed to the preparation of the manuscript. P.P. conceived and supervised the project and wrote the manuscript. C.C. and E.K contributed supporting the irradiation experiments at the Radiation Physics department.

## [Author Responsible for Statistical Analysis Name & Email Address]

Sebastián A. Barrientos

## References

1. Moore, B. D., Copeland, D. R., Ried, H. & Levy, B. Neurophysiological basis of cognitive deficits in long-term survivors of childhood cancer. Arch Neurol 49, 809–17 (1992).

2. Roman, D. D. & Sperduto, P. W. Neuropsychological effects of cranial radiation: current knowledge and future directions. Int J Radiat Oncol Biol Phys 31, 983–98 (1995).

3. Padovani, L., André, N., Constine, L. S. & Muracciole, X. Neurocognitive function after radiotherapy for paediatric brain tumours. Nat Rev Neurol 8, 578–588 (2012).

4. Makale, M. T., McDonald, C. R., Hattangadi-Gluth, J. A. & Kesari, S. Mechanisms of radiotherapy-associated cognitive disability in patients with brain tumours. Nat Rev Neurol 13, 52–64 (2017).

5. Simmons, D. A. et al. Reduced cognitive deficits after FLASH irradiation of whole mouse brain are associated with less hippocampal dendritic spine loss and neuroinflammation. Radiother Oncol 139, 4–10 (2019).

6. Pazzaglia, S., Briganti, G., Mancuso, M. & Saran, A. Neurocognitive Decline Following Radiotherapy: Mechanisms and Therapeutic Implications. Cancers (Basel) 12, (2020).

7. Rübe, C. E., Raid, S., Palm, J. & Rübe, C. Radiation-Induced Brain Injury: Age Dependency of Neurocognitive Dysfunction Following Radiotherapy. Cancers (Basel) 15, (2023).

8. Mulhern, R. K. et al. Neurocognitive consequences of risk-adapted therapy for childhood medulloblastoma. J Clin Oncol 23, 5511–5519 (2005).

9. Ivica, N., Tamté, M., Ahmed, M., Richter, U. & Petersson, P. Design of a high-density multi-channel electrode for multi-structure parallel recordings in rodents. (2014) doi:10.1109/EMBC.2014.6943611.

10. Semple, B. D., Blomgren, K., Gimlin, K., Ferriero, D. M. & Noble-Haeusslein, L. J. Brain development in rodents and humans: Identifying benchmarks of maturation and vulnerability to injury across species. Prog Neurobiol 106–107, 1–16 (2013).

11. Bressler, S. L. & Menon, V. Large-scale brain networks in cognition: emerging methods and principles. Trends Cogn Sci 14, 277–290 (2010).

12. Moran, R. J. et al. Alterations in brain connectivity underlying beta oscillations in Parkinsonism. PLoS Comput Biol 7, e1002124 (2011).

13. Santana, M. et al. Spinal Cord Stimulation Alleviates Motor Symptoms in a Primate Model of Parkinson’s disease. Neuron (84), 1–7 (2014).

14. Monje, M. & Fisher, P. G. Neurological complications following treatment of children with brain tumors. J Pediatr Rehabil Med 4, 31–6 (2011).

15. Antunes, M. & Biala, G. The novel object recognition memory: neurobiology, test procedure, and its modifications. Cogn Process 13, 93–110 (2012).

16. Clark, R. E., Zola, S. M. & Squire, L. R. Impaired Recognition Memory in Rats after Damage to the Hippocampus. Journal of Neuroscience 20, 8853–8860 (2000).

17. Vogel-Ciernia, A. & Wood, M. A. Examining object location and object recognition memory in mice. Curr Protoc Neurosci 69, 8.31.1–8.31.17 (2014).

18. Paxinos, G. & Watson, C. The rat brain in stereotaxic coordinates. Elsevier, 6th Edition, Academic Press, San Diego. (2007).

19. DeCoteau, W. E. et al. Learning-related coordination of striatal and hippocampal theta rhythms during acquisition of a procedural maze task. Proc Natl Acad Sci U S A 104, 5644– 5649 (2007).

20. Fujisawa, S. & Buzsáki, G. A 4-Hz oscillation adaptively synchronizes prefrontal, VTA and hippocampal activities. Neuron 72, 153 (2011).

21. Paul L. Nunez; Ramesh Srinivasan. Electric Fields of the Brain - The Neurophysics of EEG. (Oxford University Press, 2005).

22. Herreras, O. Local Field Potentials: Myths and Misunderstandings. Front Neural Circuits 10, 101 (2016).

23. Womelsdorf, T. & Fries, P. The role of neuronal synchronization in selective attention. Curr Opin Neurobiol 17, 154–160 (2007).

24. Colgin, L. L. et al. Frequency of gamma oscillations routes flow of information in the hippocampus. Nature 462, 353–357 (2009).

25. Wen, H. & Liu, Z. Separating Fractal and Oscillatory Components in the Power Spectrum of Neurophysiological Signal. Brain Topogr 29, 13–26 (2016).

26. Alonso, J. M. & Swadlow, H. A. Thalamus controls recurrent cortical dynamics. Nat Neurosci 18, 1703–1704 (2015).

27. Drayson, O. G., et al. A multi-institutional study to investigate the sparing effect after whole brain electron FLASH in mice: Reproducibility and temporal evolution of functional, electrophysiological, and neurogenic endpoints. bioRxiv 2024.01.25.577164 (2024) doi:10.1101/2024.01.25.577164.

28. Wu, M. Y. et al. Cranial irradiation impairs intrinsic excitability and synaptic plasticity of hippocampal CA1 pyramidal neurons with implications for cognitive function. Neural Regen Res 17, 2253–2259 (2022).

29. Kahalley, L. S. et al. Slower processing speed after treatment for pediatric brain tumor and acute lymphoblastic leukemia. Psychooncology 22, 1979–1986 (2013).

30. Jacobson, L. A., Mahone, E. M., Yeates, K. O. & Ris, M. D. Processing speed in children treated for brain tumors: effects of radiation therapy and age. Child Neuropsychology 25, 217–231 (2019).

31. Irestorm, E., Ora, I., Linge, H. & Tonning Olsson, I. Cognitive Fatigue and Processing Speed in Children Treated for Brain Tumours. Journal of the International Neuropsychological Society 27, 865–874 (2021).

32. Montay-Gruel, P. et al. Long-term neurocognitive benefits of FLASH radiotherapy driven by reduced reactive oxygen species. Proc Natl Acad Sci U S A 166, 10943–10951 (2019).

33. Bevins, R. A. & Besheer, J. Object recognition in rats and mice: a one-trial non-matching-to-sample learning task to study ‘recognition memory’. Nat Protoc 1, 1306–1311 (2006).

34. Dix, S. L. & Aggleton, J. P. Extending the spontaneous preference test of recognition: Evidence of object-location and object-context recognition. Behavioural Brain Research 99, 191–200 (1999).

35. Mathis, A. et al. DeepLabCut: markerless pose estimation of user-defined body parts with deep learning. Nat Neurosci 21, 1281–1289 (2018).

36. Hampson, R. E., Heyser, C. J. & Deadwyler, S. A. Hippocampal cell firing correlates of delayed-match-to-sample performance in the rat. Behavioral neuroscience 107, 715–739 (1993).

37. Bokil, H., Andrews, P., Kulkarni, J. E., Mehta, S. & Mitra, P. P. Chronux: A Platform for Analyzing Neural Signals. J Neurosci Methods 192, 146 (2010).

38. Fukuda, A. et al. Age-dependent sensitivity of the developing brain to irradiation is correlated with the number and vulnerability of progenitor cells. J Neurochem 92, 569–84 (2005).

39. Naylor, A. S. et al. Voluntary running rescues adult hippocampal neurogenesis after irradiation of the young mouse brain. Proc Natl Acad Sci U S A 105, 14632–7 (2008).

40. Blomstrand, M., Kalm, M., Grandér, R., Björk-Eriksson, T. & Blomgren, K. Different reactions to irradiation in the juvenile and adult hippocampus. Int J Radiat Biol 90, 807–15 (2014).

41. Zanni, G. et al. Irradiation of the Juvenile Brain Provokes a Shift from Long-Term Potentiation to Long-Term Depression. Dev Neurosci 37, 263–72 (2015).

42. Eichenbaum, H. Hippocampus: Cognitive Processes and Neural Representations that Underlie Declarative Memory. Neuron 44, 109–120 (2004).

43. Haber, S. N. Corticostriatal circuitry. Dialogues Clin Neurosci 18, 7 (2016).

44. Censoni, L. et al. Verification of multi-structure targeting in chronic microelectrode brain recordings from CT scans. J Neurosci Methods 382, (2022).

