## Supplementary Figures for "Aberrant Neuronal Synchronization Associated with Cognitive Deficits in a Rodent Model of Childhood Cranial Irradiation"

### Supp 1

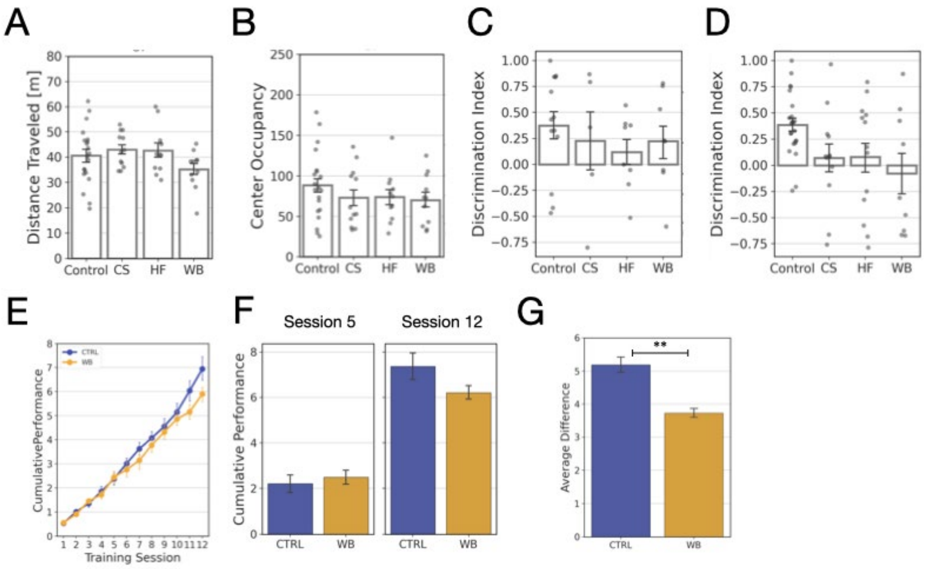


**Supplementary Figure 1 – Behavioral tests in CTRL and WB rats**

**A)** Average Distance Travelled per animal during exploration of the OF. **B)** Average Center Occupancy per animal during exploration of the OF. **A-B)** $N_{CTRL\text{-}CS\text{-}HF\text{-}WB}$ = [20, 12, 11, 11]. **C)** Average Discrimination Index per recorded animal during the NOR test. $N_{CTRL\text{-}CS\text{-}HF\text{-}WB}$ = [13, 5, 8, 8]. **D)** Average Discrimination Index per recorded animal during the OLR test. $N_{CTRL\text{-}CS\text{-}HF\text{-}WB}$ = [23, 12, 13, 9]. **A-D)** Note that although a Kruskal-Wallis test showed no significance between the groups, significant group-specific differences found against Control are specified in the main text. **E)** Cumulative performance during the training period of the cued NMTS task for CTRL and WB rats. **F)** Cumulative performance at Session 5 (left, $N_{CTRL\text{-}WB}$= [10, 7]) and Session 12 (right, $N_{CTRL\text{-}WB}$= [6, 8]) for CTRL and WB rats. **G)** Average difference between Session 12 and Session 5 for both treatment groups. Bars and Markers indicate the mean across recorded animals and error bars indicate SEM. Significance was evaluated by Bootstrapping test between Control and WB (** p-value < 0.001)

#### Supp 2

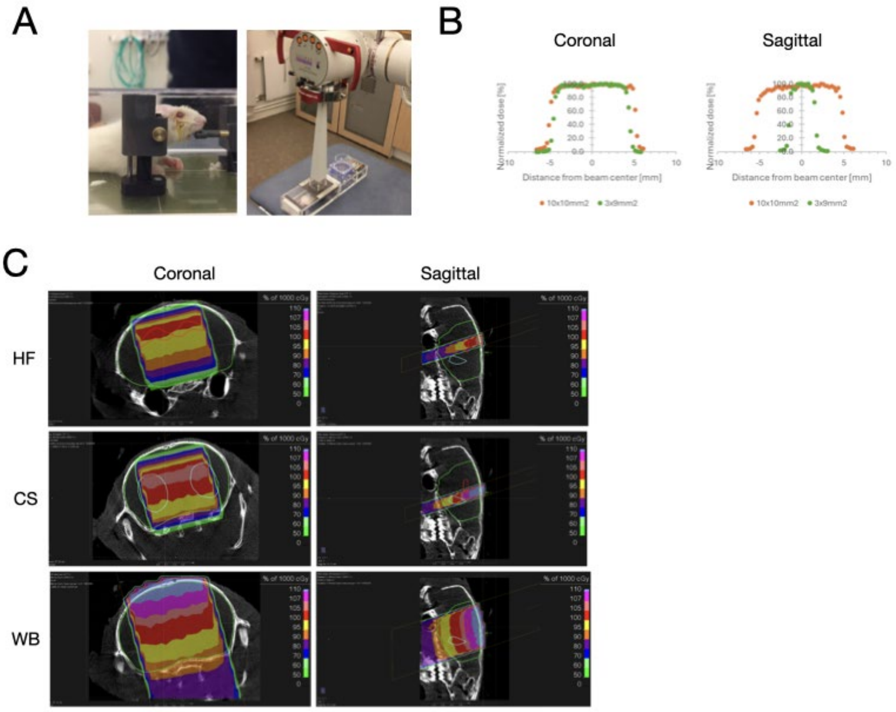


**Supplementary Figure 2 - Irradiation Design**

**A)** Example of a rat pup (21 post-natal days) subjected to cranial irradiation in prone position in a ventilated chamber. **B)** Measured dose profiles for the different apertures. **C)** Simulation of the absorbed dose distribution for the three different target regions HF, CS and WB**.**

#### Supp 3

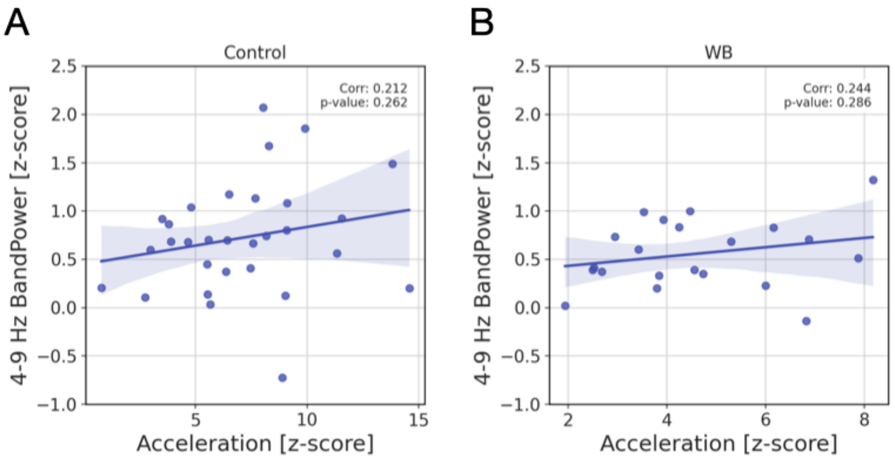


**Supplementary Figure 3 - Correlation between normalized theta band power and acceleration recorded by head-mounted accelerometers.**

**A)** Normalized accelerometer data (z-score) collected during the full running period compared against the corresponding Normalized 4-9 Hz Bandpower (z-score) for Control [N = 30] and, **B)** WB rats [N = 21]. **A-B)** Each dot represents the average values across grouped datapoints (Recording, Outcome, Treatment and Epoch). The diagonal line depicts the linear regression with the shaded area indicating the confidence interval. Note that a similar non-significant trend of increased theta with higher motility was observed in both treatment groups (increased theta power during locomotion has indeed been reported in previous studies [43] ). Pearson Correlation coefficients and p-value are displayed in the plots.
